# Cerebellar Purkinje cell microcircuits are essential for tremor

**DOI:** 10.1101/770321

**Authors:** Amanda M Brown, Joshua J White, Meike E van der Heijden, Tao Lin, Roy V Sillitoe

## Abstract

Tremor is currently ranked as the most common movement disorder. The brain regions and neural signals that initiate the debilitating shakiness of different body parts remain unclear. Here, we found that genetically silencing cerebellar Purkinje cell activity blocked tremor in mice that were given the tremorgenic drug harmaline. We show in awake behaving mice that the onset of tremor is coincident with rhythmic Purkinje cell firing, which alters the output of their target cerebellar nuclei cells. We mimic the tremorgenic action of the drug with optogenetics and present evidence that highly patterned Purkinje cell activity drives a powerful tremor in otherwise normal mice. Modulating the altered activity with deep brain stimulation directed to the Purkinje cell output in the cerebellar nuclei reduced tremor in freely moving mice. Together, the data implicate Purkinje cell connectivity as a neural substrate for tremor and a gateway for signals that mediate the disease.

## Introduction

Tremors are uncontrollable muscle oscillations that result in rhythmic shaking of the affected body parts. Tremor occurs in healthy individuals at baseline, which is known as physiological tremor (*1*). However, tremor also occurs as a movement disorder when its amplitude becomes severe enough to disrupt daily activities (*2*). Tremor can constitute independent diseases, such as in essential tremor, the most common tremor disease (*3, 4*). It can also be co-morbid with other brain disorders such as Parkinson’s disease (*5*), dystonia (*6, 7*), ataxia (*8*), or epilepsy (*9*). Additionally, tremor can be a negative consequence of a growing list of common prescription drugs, toxins, or neurological insults (*10, 11*). While there are a great number of diseases, disorders, and chemicals that are associated with tremor, the neural origins of tremor are largely not understood and they are especially unclear in the most common tremor diseases (*5, 12*).

There is good evidence implicating dysfunctional cerebello-thalamo-cortical circuits in tremor. The cerebellar receiving areas of the thalamus such as the ventral intermediate nucleus (VIM) and the ventral anterolateral nucleus (VAL) are preferred targets for thalamotomy and deep brain stimulation (DBS) in the treatment of essential tremor (*13*). Local field potentials and spike activity recorded from these brain areas in humans experiencing bouts of tremor correlate with the frequency of oscillation in the affected body parts (*12, 14*). However, it is unclear where in the brain this abnormal activity originates (*12*). Functional magnetic resonance imaging (fMRI) studies in humans with essential tremor reported abnormal levels of activity in the cerebellum (*15*). Compellingly, when brain activity of individuals with tremor disorders is compared between periods of mimed tremor and true epochs of tremor, the only area of the brain with significantly different patterns of activity is the cerebellum (*16*). Yet, if and how the cerebellum could provide a major contribution to either generating the tremor–therefore, acting as an origin of the signal–or mediating the transfer of an existing tremor signal, has not been elucidated. Further, the respective role that individual cerebellar cell types may have *in vivo* in the behaving animal during the production or propagation of tremor-related neural activity is unknown.

Abnormalities in different cerebellar cell types, particularly the Purkinje cells, have been associated with tremor. However, it is unclear whether these neurons are directly responsible for generating or propagating tremor (*17*). Likely, the positioning of each cell type within the local cerebellar circuitry, as well as the motor circuit as a whole, influences how each one influences tremor (*18*). The cerebellar cortex has a canonical and repeating architecture throughout all of its lobules and is comprised of the Purkinje cells at the center of a microcircuit that integrates information from five major classes of excitatory and inhibitory interneurons. Inputs to the cerebellum include mossy fibers and climbing fibers, with the latter originating in the inferior olive where it sends powerful excitatory inputs directly onto the Purkinje cell dendrites. Purkinje cells project out of the cerebellar cortex to make inhibitory synapses onto the cerebellar nuclei neurons. The cerebellar nuclei provide the final output of the cerebellum, representing the culmination of all cerebellar inputs and computations therein. Therefore, current views consider the cerebellar nuclei signals as a link between the cerebellum and the rest of the brain and spinal cord and Purkinje cell activity as the computational center that shapes these signals (*19*) (**Fig. 1a-c**).

**Fig. 1:**
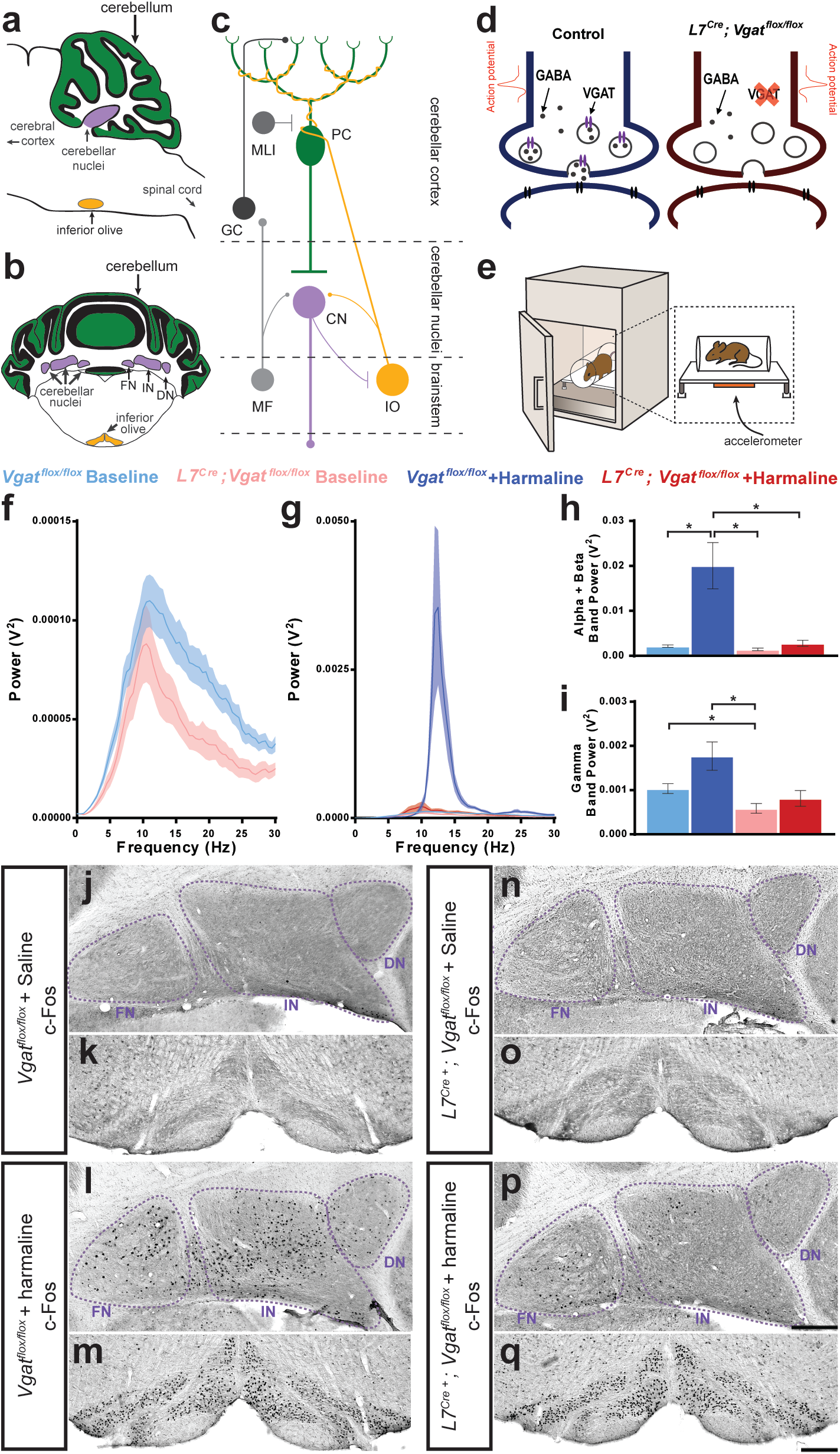
Purkinje cell neurotransmission is necessary for producing baseline physiological tremor and pathological tremor. (**a**) Representation of a sagittal section of a mouse cerebellum indicating its spatial relationship to other landmarks of the central nervous system. Green region = Purkinje cell dendrites and molecular layer, purple region = cerebellar nuclei, yellow region = inferior olive. (**b**) Representation of a coronal section through a mouse cerebellum where the fastigial nucleus (FN), interposed nucleus (IN), and dentate nucleus (DN) are all visible. Green region = Purkinje cell dendrites and molecular layer, purple regions = cerebellar nuclei, yellow regions = inferior olive. (**c**) Representation of a simplified cerebellar circuit including a Purkinje cell (PC, green), cerebellar nuclei (CN, purple), inferior olive (IO, yellow), mossy fibers (MF), granule cell (GC), and molecular layer interneuron (MLI). Large circles = cell bodies, small circle terminals = excitatory synapses, flat terminals = inhibitory synapses. (**d**) Representation of the result of genetic manipulation in *L7^Cre^;Vgat^flox/flox^* mice. Control Purkinje cell synapse depicted in blue on left, *L7^Cre^;Vgat^flox/flox^* Purkinje cell synapse depicted in red on right. Large open circles = vesicles. Small filled circles = GABA. Purple ellipse pairs = VGAT. Bright red action potential cartoon represents an action potential reaching the synapse and triggering the fusion of vesicles to the presynaptic membrane and release of the vesicles’ contents, such as GABA, onto receptors in the postsynaptic membrane (black ellipse pairs). GABA is released from Purkinje cells during fast neurotransmission in *Vgat^flox/flox^* control mice, but not in *L7^Cre^;Vgat^flox/flox^* mice. (**e**) Representation of a commercial tremor monitor. Inset = dotted rectangle. Accelerometer = orange rectangle. (**f-g**) Solid line = mean. Shaded region = standard error of the mean (SEM). Legend above. (**f**) Mice lacking Purkinje cell GABA neurotransmission had lower baseline physiological tremor compared to control animals. Control N = 18, mutant N = 12. (**g**) While control animals exhibited the typical robust tremor after harmaline administration (N = 16), *L7^Cre^;Vgat^flox/flox^* animals had no significant increase in tremor in response to the drug (N = 13). The baseline data from (**f**) is repeated on this graph for scale. (**h**) Summed tremor power within the alpha and beta bands. Legend above. (**i**) Summed tremor power within the gamma band. Legend above. (**j-q**) c-Fos expression in the cerebellar nuclei (**j, l n, p**) and inferior olive (**k, m, o, q**) after saline (**j-k, n-o**) or harmaline (**l-m, p-q**) administration. Cerebellar nuclei scale = 250µm. Inferior olive scale = 250µm.

Accordingly, there is compelling, albeit indirect, evidence pointing to a role for abnormal, reduced, or the loss of Purkinje cell to cerebellar nuclei communication as a key neural substrate for tremor (*20*). Additionally, studies have found abnormalities in cells directly upstream (*21*) as well as in the cerebellar nuclei directly downstream (*22*) of Purkinje cells to be associated with tremor. While Purkinje cell loss and degenerative Purkinje cell morphology have been noted in some types of tremor, these hallmarks are not found across all tremor conditions (*11, 23*). Indeed, with the many varied potential causes and diseases associated with tremor, there may be equally as many potential biological substrates of tremor. In the face of this problem, we have sought to determine whether cerebellar Purkinje cells have a direct role in tremor generation and, if so, whether there are electrophysiological abnormalities in the cerebellum that dictate the tremor state.

## Results

### Lack of Purkinje cell GABA neurotransmission does not induce pathological tremor

Previous human pathology studies of essential tremor raised the possibility that loss or reduction of Purkinje cell signaling causes tremor (*22, 24*). In order to address whether Purkinje cells have a role in the production and propagation of tremor signals, we first tested whether removing Purkinje cell to cerebellar nuclei neurotransmission triggers tremor. To accomplish this, we used an *L7^Cre^;Vgat^flox/flox^* conditional genetic approach to delete the vesicular GABA transporter (VGAT) from Purkinje cells (*25, 26*). The result of this manipulation is that Purkinje cells can still fire action potentials, but they can no longer communicate with their downstream partners using GABA neurotransmission, which ultimately results in silencing Purkinje cell output (**Fig. 1d**). The anatomical fidelity of the cerebellar circuit is maintained despite this manipulation, though ataxia is present (*27*)(**Movie S1**). A possible additional outcome in mice with silenced Purkinje cells would be a tremor phenotype. However, we found that mice without Purkinje cell output did not have an enhanced tremor phenotype. Instead, the lack of Purkinje cell GABA neurotransmission resulted in a lower than normal baseline of physiological tremor (**Fig. 1e-f, 1i, Table 1**. *Vgat^flox/flox^* (referred to from here on as control) *baseline N = 18, L7^Cre^;Vgat^flox/flox^* (referred to from here on as mutant) *baseline N = 12*). These data suggest that Purkinje cell activity may have a role in establishing the normal level of physiological tremor but the lack of Purkinje cell activity alone is not a sufficient change in the circuit to result in pathological levels of tremor.

### Lack of Purkinje cell neurotransmission reduces harmaline tremor

Since losing Purkinje cell activity did not produce tremor, we next sought to determine the role of Purkinje cells in the context of a potentially greater tremor circuit. For this, we administered harmaline, a beta-carboline alkaloid compound that causes an 11-14 Hz tremor in mice and 8-16 Hz tremor in multiple species, including humans (*28*). Harmaline affects many types of receptors, ion channels, and gap junctions that are found throughout the nervous system, and therefore likely affects the activity of multiple cell populations in the brain (*29*). However, a rich history of research in slice (*30*), decerebrate (*31, 32*), and anesthetized (*30, 32*) preparations have indicated that harmaline induced tremor involves synchronous rhythmic firing in the inferior olive. If this is the case, then one would postulate that Purkinje cells must be involved in the production of the tremor response. The influence of Purkinje cells during this process has remained unclear due to complicated results in genetic models, diverse circuit manipulation techniques, a confounding lack of cell type specificity, or the presence of degeneration that interferes with interpretation of neuronal function (*32–34*). We found that mice lacking Purkinje cell signaling did not have a significantly increased tremor after harmaline administration compared to control animals, which displayed the typical robust tremor in response to the drug (**Fig. 1e, 1g-h, Table 1, Movie S2**. *Control + harmaline N = 16, mutant + harmaline N = 13*). We next asked how key nodes in the cerebellar system collectively respond to changes in circuit activity after harmaline is provided. Harmaline administration resulted in robust activation of the early activation transcription factor c-Fos in both the cerebellar nuclei and inferior olive of control animals (**Fig. 1j-m**). However, in mice with silenced Purkinje cell output, the cerebellar nuclei were largely spared the abnormal activation of c-Fos (**Fig. 1n, 1p**). This is despite having similar activation of the inferior olive (**Fig. 1o, 1q**). These data suggest that Purkinje cell signaling is required for propagating the neural activity that drives harmaline induced tremor, and that Purkinje cell to cerebellar nuclei communication is an essential synapse for promoting tremor behavior.

### Harmaline causes Purkinje cells to develop a bursting pattern of activity in awake behaving mice

As our data pointed to Purkinje cell activity as a primary factor in mediating tremor, we sought to define the underlying Purkinje cell activity that occurs during tremor. We performed single unit extracellular recordings of Purkinje cells in awake head-fixed mice both before and during tremor that was triggered by the acute effects of harmaline (**Fig. 2a-c**). This allowed us to record and quantify both the Purkinje cells’ simple spikes, which are both intrinsically generated and modulated by mossy fiber inputs, as well as complex spikes which are generated via the climbing fiber input (**Fig. 1c, 2c**). In both control and mutant animals, Purkinje cell simple spike activity developed a dramatic bursting pattern during tremor (**Fig. 2d-g**). When we quantified the spike properties of these cells, we found that Purkinje cell simple spike activity had a significantly decreased frequency and significantly increased coefficient of variance (CV) and CV2 during tremor (*35*) (**Fig. 2h-j, Table 2.** *Control baseline N = 4, n = 18. Control + harmaline N = 6, n = 14. Mutant baseline N = 4, n = 15. Mutant + harmaline N = 5, n = 12*). CV is a measure of irregularity of interspike intervals over the entire recording of the cell, and therefore a higher CV value indicates a greater overall bursting quality of cell activity. Meanwhile, CV2 is a measure of irregularity of directly adjacent interspike intervals, and therefore a higher CV2 value indicates a more erratic and unpredictable quality of a spike train. However, CV2 can also be elevated when there is an overall bursting quality of cell activity, especially when the number of spikes within a burst or the overall firing rate is low, as we have shown here (**Fig. 2e, 2g-h**) (*35*). Therefore, simple spike activity predominantly decreases in frequency and increases in “burstiness” after harmaline administration.

**Fig. 2:**
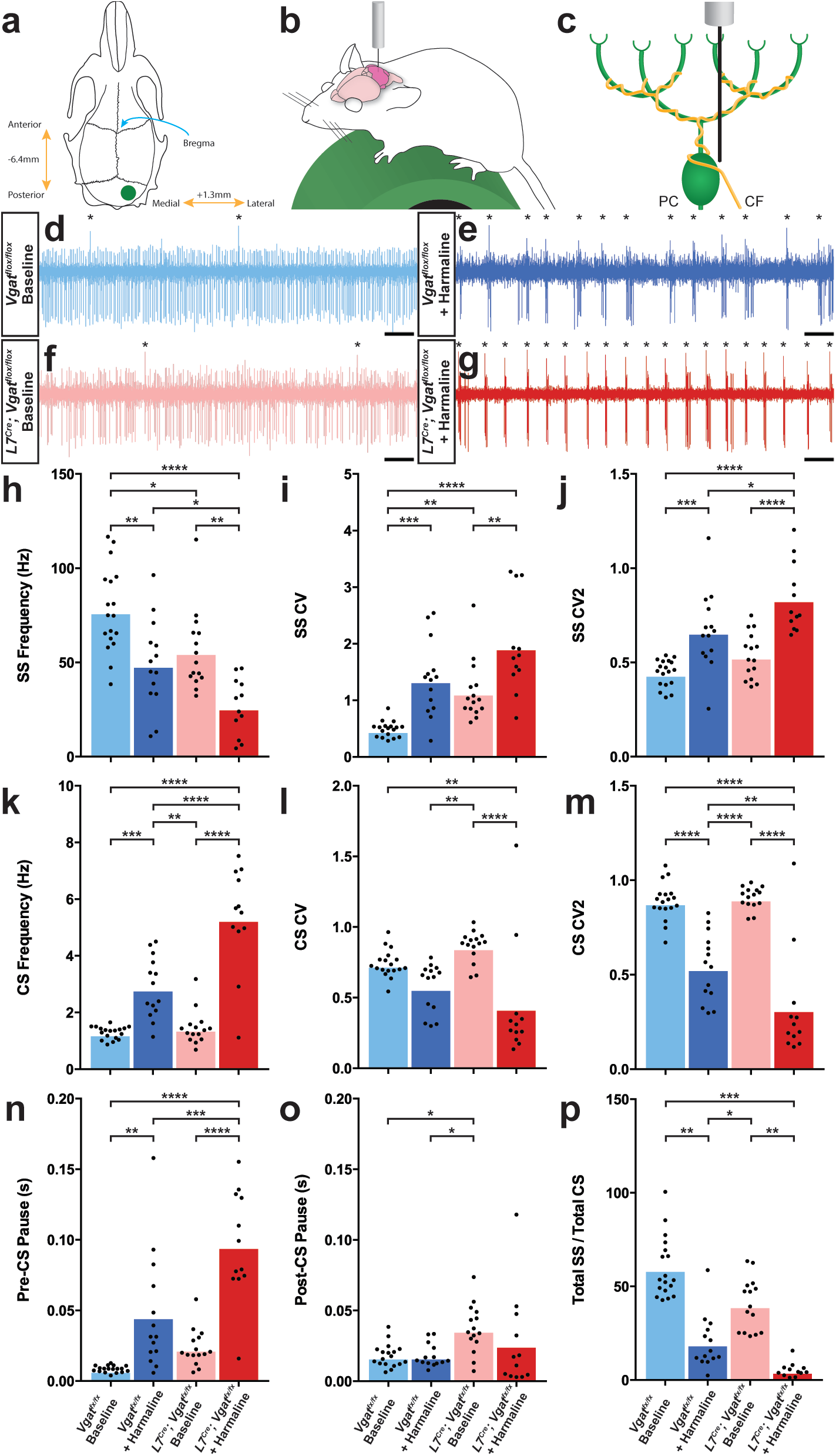
Purkinje cell firing patterns are significantly altered after harmaline administration. (**a**) Representation of craniotomy site using skull landmarks. Craniotomy (green circle) for awake head-fixed neural recordings was made −6.4mm from Bregma (blue arrow) and 1.3mm lateral from midline. (**b**) Representation of awake head-fixed recordings. Mice were allowed to stand on a green foam wheel (green cylinder) during recordings. (**c**) Representation of extracellular recordings of Purkinje cells (PC) which allowed recordings of simple spikes and complex spikes, which are triggered by the climbing fiber (CF). (**d-g**) Example raw trace recordings of Purkinje cells. Complex spikes are indicated with asterisks. Scale = 500ms. (**d**) Purkinje cell from a control animal. (**e**) Purkinje cell from a control animal during tremor after harmaline administration. (**f**) Purkinje cell from a mutant animal. (**g**) Purkinje cell from a mutant animal after harmaline administration (tremor not present). (**h-j**) Quantification of Purkinje cell simple spike firing properties including frequency (**h)**, CV (**i**), and CV2 (**j**). (**k-m**) Quantification of Purkinje cell complex spike firing properties including frequency (**k)**, CV (**l**), and CV2 (**m**). (**n-p**) Quantification of Purkinje cell simple spike and complex spike relationship including pre complex spike pause duration (**n)**, post complex spike pause duration (**o**), and total simple spike to complex spike ratio (**p**).

Complex spike properties changed in the opposite direction, wherein frequency was significantly increased and CV2 significantly decreased, while CV was not significantly altered in control animals (**Fig. 2k-m, Table 2.** *Control baseline N = 4, n = 18. Control + harmaline N = 6, n = 14. Mutant baseline N = 4, n = 15. Mutant + harmaline N = 5, n = 12*). Bursts of activity tended to be led by a complex spike and preceded by a pause in activity (**Fig. 2e, 2g**). This is evidenced by a significantly greater duration of inter-spike interval (ISI) before each complex spike and no change in duration after each complex spike after harmaline administration (**Fig. 2n-o, Table 2.** *Control baseline N = 4, n = 18. Control + harmaline N = 6, n = 14. Mutant baseline N = 4, n = 15. Mutant + harmaline N = 5, n = 12.*). Therefore, complex spike timing properties also apply to burst timing properties. Since simple spike frequency decreased and the complex spike frequency increased, this resulted in an overall decrease in simple spike to complex spike ratio during tremor (**Fig. 2p, Table 2.** *Control baseline N = 4, n = 18. Control + harmaline N = 6, n = 14. Mutant baseline N = 4, n = 15. Mutant + harmaline N = 5, n = 12.*). Together, these data indicate that Purkinje cell output is defined by an overall lower simple spike firing rate and a steady pattern of simple spike bursts after harmaline administration in an awake behaving condition. These bursts are flanked by complex spikes that are increased in frequency and regularity. In all quantified measures of spike activity, a similar directionality of change occurred in the mutant Purkinje cells of mice with silenced Purkinje cell output compared to those in the controls. These electrophysiological data suggest that if the circuits that carry tremor related neural activity eventually innervate the Purkinje cells, then the fidelity of the pathways that transfer the signals is equivalent in mutants and controls since both genotypes of mice had similar *in vivo* neuronal responses to the drug (**Fig. 2h-p**).

### Purkinje cell neurotransmission is necessary for propagating harmaline-induced burst activity

We next tested what affect the abnormal Purkinje cell activity has on the downstream neurons in the cerebellar nuclei because these cells provide the major output of the cerebellum. We performed single cell extracellular recordings of cerebellar nuclei cells in awake head-fixed mice both before and during tremor caused by harmaline (**Fig. 3a**). We found significantly different responses in the cerebellar nuclei of control mice compared to those lacking Purkinje cell signaling (**Fig. 3b-e**). Cerebellar nuclei cells in control animals experiencing tremor had a significant and more predominant bursting pattern as measured by CV with no significant change in firing frequency or CV2 from baseline (**Fig. 3f-h, Table 3**). However, cerebellar nuclei cells in animals lacking Purkinje cell GABA neurotransmission – which results in little to no tremor after harmaline administration – had no change from baseline in any of our measures of cerebellar nuclei spike properties (**Fig. 3f-h, Table 3.** *Control baseline N = 5, n = 19. Control + harmaline N = 3, n = 14. Mutant baseline N = 6, n = 18. Mutant + harmaline N = 4, n = 11.*). Moreover, the burst activity of the cerebellar output appears highly rhythmic. These data suggest that *in vivo* circuit alterations that promote abnormal burst activity in the cerebellar nuclei, with Purkinje cell signals as one major source, could drive the core behavioral features of tremor in behaving mice.

**Fig. 3:**
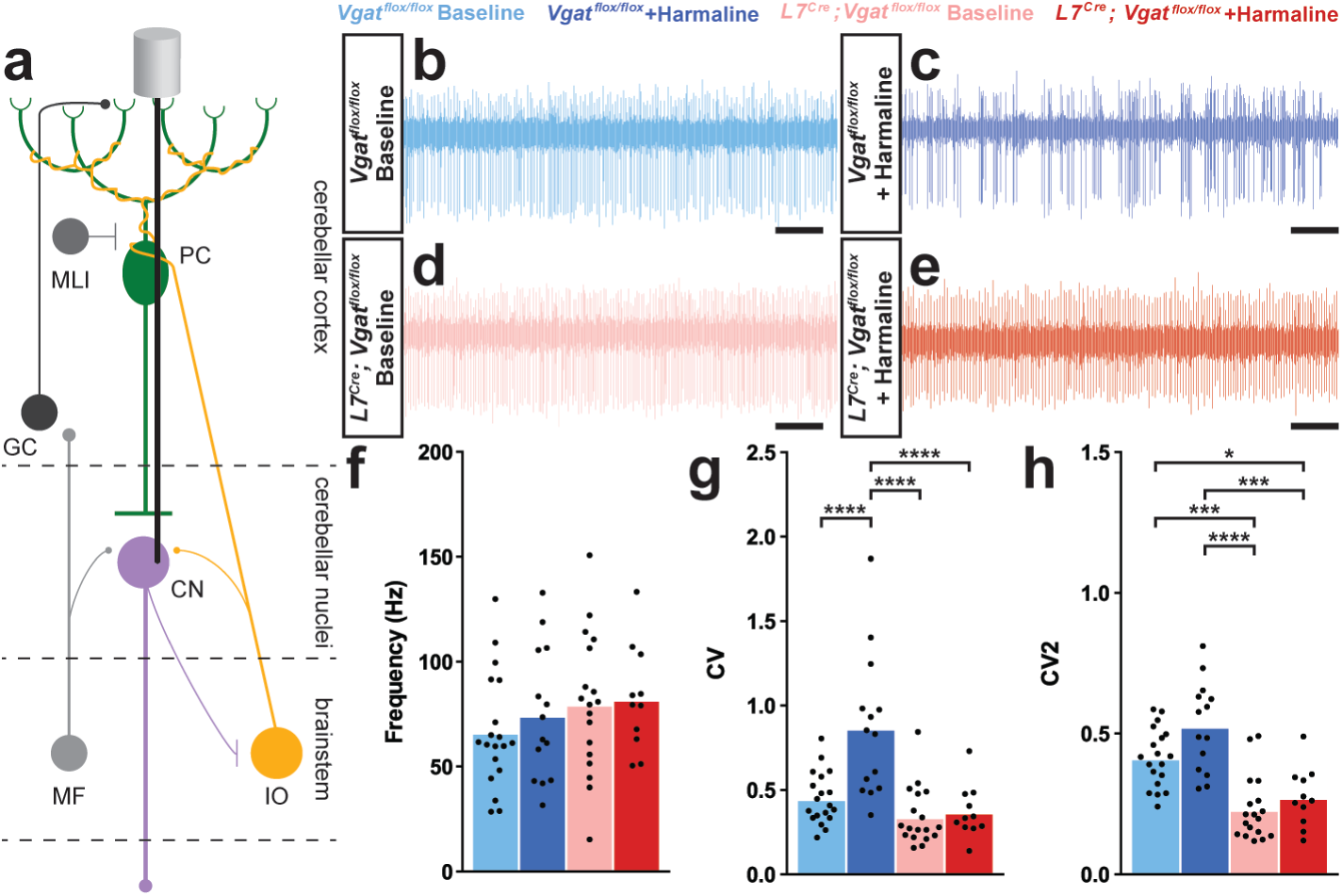
A burst pattern of cerebellar nuclei activity is associated with the tremor phenotype. (**a**) Representation of an extracellular recording of the cerebellar nuclei. (**b-e**) Example raw trace from recordings of cerebellar nuclei cells. Scale = 250ms. (**b**) Cerebellar nuclei cell from a control animal. (**c**) Cerebellar nuclei cell from a control animal during tremor, after harmaline administration. (**d**) Cerebellar nuclei cell from a mutant animal. (**e**) Cerebellar nuclei cell from a mutant animal after harmaline administration (tremor not present). (**f**) Quantification of population cerebellar nuclei firing frequency. (**g**) Quantification of population cerebellar nuclei CV. (**h**) Quantification of population cerebellar nuclei CV2.

### Rhythmic bursting activity in the cerebellar nuclei produces tremor across a range of frequencies

The genetic manipulation of Purkinje cells in control and harmaline treated mice argues that, whereas losing Purkinje cell activity is not a driver for tremor, changing their pattern and mode of interaction with the cerebellar nuclei might be. We subsequently tested whether recreating the abnormal cerebellar activity found in control harmaline treated mice is sufficient for producing tremor. We implanted optical fibers bilaterally into the interposed nuclei of *L7^Cre^;ROSA26^flox-stop-EYFP-ChR2^* mice in which the opsin is only expressed in Purkinje cells. We also implanted EMG electrodes into the gastrocnemius of the left hind limb to measure tremor with a particular emphasis on examining muscle activity that occurs during movements (**Fig. 4a-c**). We stimulated Purkinje cell terminals in the cerebellar nuclei with sinusoidal pulses of light at 1-20 Hz to induce different frequencies of bursting activity of the cerebellar nuclei (**Fig. 4d-j**). We found that inducing bursting patterns of cerebellar nuclei activity resulted in tremor (**Fig. 4k-l, Movie S3**). Tremor could be elicited at all frequencies tested and was visible by eye and detectable in the EMG trace (**Fig. 4k-p**). The predominant frequency of tremor elicited matched the frequency of optogenetic stimulation and was only present during stimulation (**Fig. 4m-o, Table 4.** *Fig. 4n: N = 7*). Interestingly, the optogenetically-elicited tremor was not uniform in severity across frequencies, with the maximum power occurring during tremor frequencies around 10 Hz and 20 Hz (**Fig. 4o-p**). These data show that Purkinje-cell-induced bursting of the cerebellar nuclei is capable of generating a phenotypically obvious tremor at a wide range of frequencies. However, the data also indicates that there may be an ideal band of frequencies and harmonics at which the cerebellum instigates the strongest responses in the muscle, which ultimately execute the tremor behavior.

**Fig. 4:**
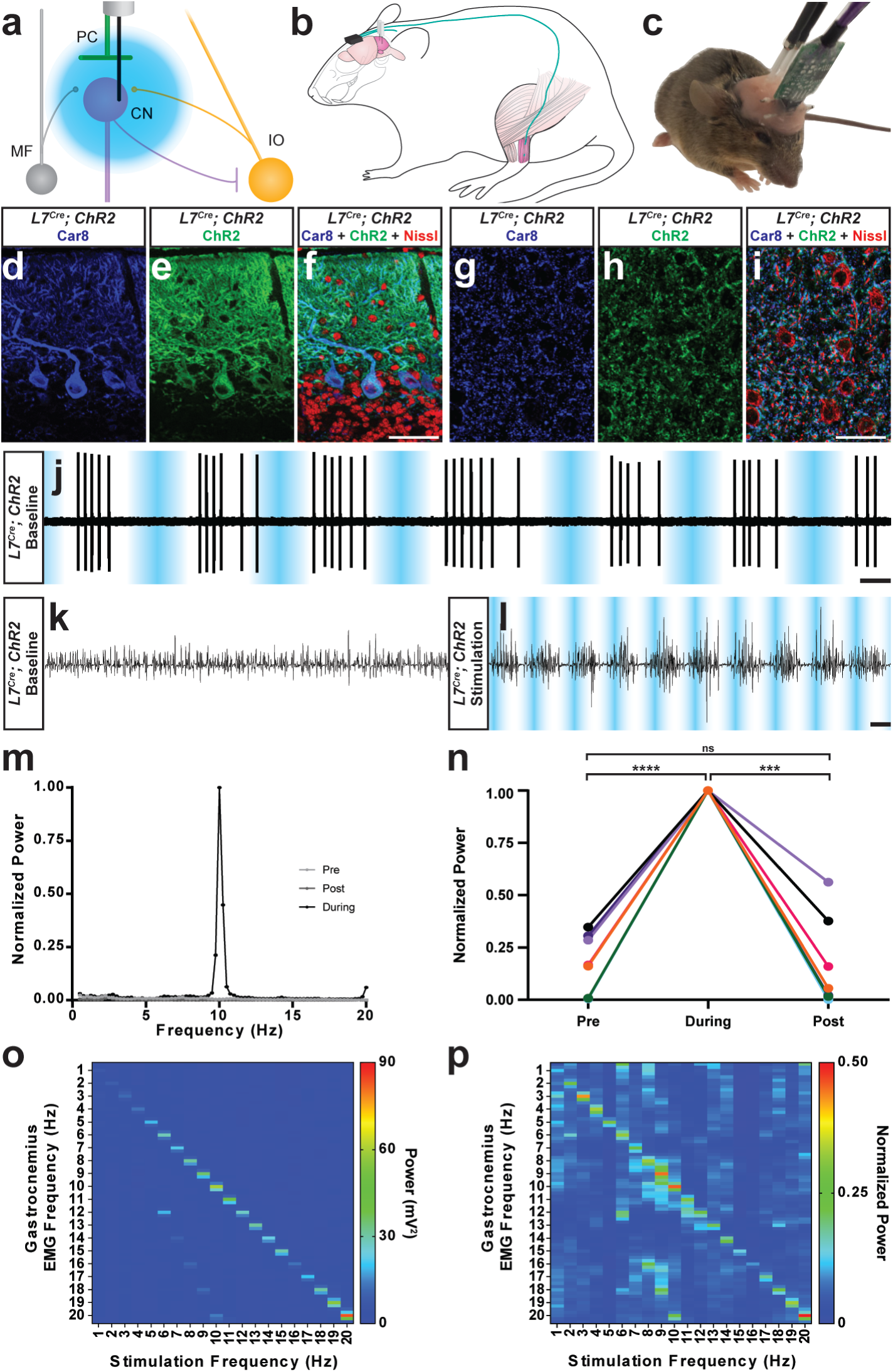
The burst pattern of cerebellar nuclei activity is sufficient to produce tremor at multiple frequencies. (**a**) Representation of an optical fiber in the cerebellar nuclei. (**b**) Representation of EMG and optical fiber implant strategy. EMG electrodes are implanted into the gastrocnemius muscle (pink) of the left hindlimb and electrode wire (teal) is fed under the skin to a connector (black) fixed to the skull. An additional wire is fed from the connector to under the skin of the neck region as a ground. Two optical fibers (white) are implanted bilaterally targeting the interposed nucleus. (**c**) Image of a mouse during an EMG recording. A preamplifier (green) is placed in the connector and tethered to a passive commutator (not pictured). Fiber patch cables (black cables) are connected to the implanted optical fibers. (**d-f**) Triple-stained fluorescent micrographs of the cerebellar cortex. Scale = 50µm. (**d**) CAR8 protein in Purkinje cells stained blue. (**e**) Membrane-bound ChR2 labeled with green fluorescent protein (GFP) stained green. (**f**) Composite of CAR8 (blue), ChR2 (green), and Nissl to label all cells (red). (**g-i**) Triple-stained fluorescent micrograph of cerebellar nuclei cells. Scale = 50µm. (**g**) CAR8 protein in Purkinje cell terminals in the cerebellar nuclei (blue). (**h**) Membrane-bound ChR2 labeled with GFP (green). (**d**) Composite of CAR8 (blue), ChR2 (green), and Nissl to label all cells (red). (**j**) Example extracellular recording from a cerebellar nuclei cell during ChR2 stimulation of surrounding Purkinje cell terminals (blue bars). Scale = 250ms. (**k-l**) Example raw EMG traces from a *L7^Cre^;ROSA26^flox-stop-ChR2-EYFP^* mouse. Scale = 50ms. (**k**) Baseline EMG trace before optogenetic stimulation. (**l**) EMG trace during tremor caused by optogenetic stimulation. Stimulation periods indicated by overlaid blue bars. (**m**) Example power spectrum analysis from an animal receiving optogenetic stimulation at 10 Hz, normalized to peak tremor power. Pre = pre-stimulation period (baseline). Post = post-stimulation period. During = during stimulation period. (**n**) EMG power at 10 Hz during 10 Hz stimulation for all mice tested, normalized to each individual’s overall maximum power in the pre, during, and post stimulation periods. Pre vs during stimulation period: P < 0.0001. During vs post stimulation period: P = 0.0001. Pre vs post stimulation period: P = 0.9796, not significant (ns). N = 7. (**o**) Heat plot showing population average elicited EMG power for each optogenetic stimulation frequency tested. Heat scale = 0 to 90mV^2^. (**p**) Heat plot showing population average of power normalized to individual peaks for each optogenetic stimulation frequency tested. Heat scale = 0 to 0.5.

### DBS directed to the cerebellar nuclei reduces tremor severity

Finally, because the data suggested that erroneous Purkinje cell to cerebellar nuclei communication was critical for the production of tremor, we aimed to correct this communication using deep brain stimulation (DBS) in order to treat ongoing tremor. We first designed and built an open tremor monitor setup that would allow DBS cables to be connected to external equipment (**Fig. 5a**) (*30*). We then devised a closed-loop DBS protocol that would constantly monitor the tremor behavior of the mouse and only trigger therapeutic stimulation during periods wherein the power of tremor was above a set threshold based on the animal’s baseline physiological tremor recording (**Fig. 5b**). DBS was directed to the interposed (middle) nuclei because they provide substantial output directly to the thalamus, a brain region linked to tremor, and since this cerebellar nucleus is critical for ongoing motion, a physiological state that is particularly sensitive to tremors that involve the cerebellum. To start, we first allowed the animals to acclimate to the tremor monitor setup and then made baseline tremor recordings (**Fig. 5c**). We then administered harmaline to induce tremor and allowed 15 minutes for the tremor to develop before moving to the DBS phase of the experiment (**Fig. 5d**). Robust tremor was elicited by harmaline in implanted animals similar to our previous results (**Fig. 5e, 1f**). Closed-loop DBS was able to reduce tremor severity every time the threshold was crossed, was automatically shut off at levels below threshold, and then was triggered again on simultaneous bouts throughout the paradigm (**Fig. 5f-g**). We quantified tremor in short 80s windows of time towards the end of the baseline period, after tremor had developed, and directly after activating stimulation. We normalized to the maximum power over the three time periods and found that tremor severity peaked during the harmaline period before stimulation was initiated and that tremor severity was decreased to levels that were not significantly different from baseline when closed-loop DBS was activated (**Fig. 5h-i, Table 5.** *N = 4*). These data show that therapeutic neuromodulation of cerebellar nuclei activity reduces tremor severity.

**Fig. 5:**
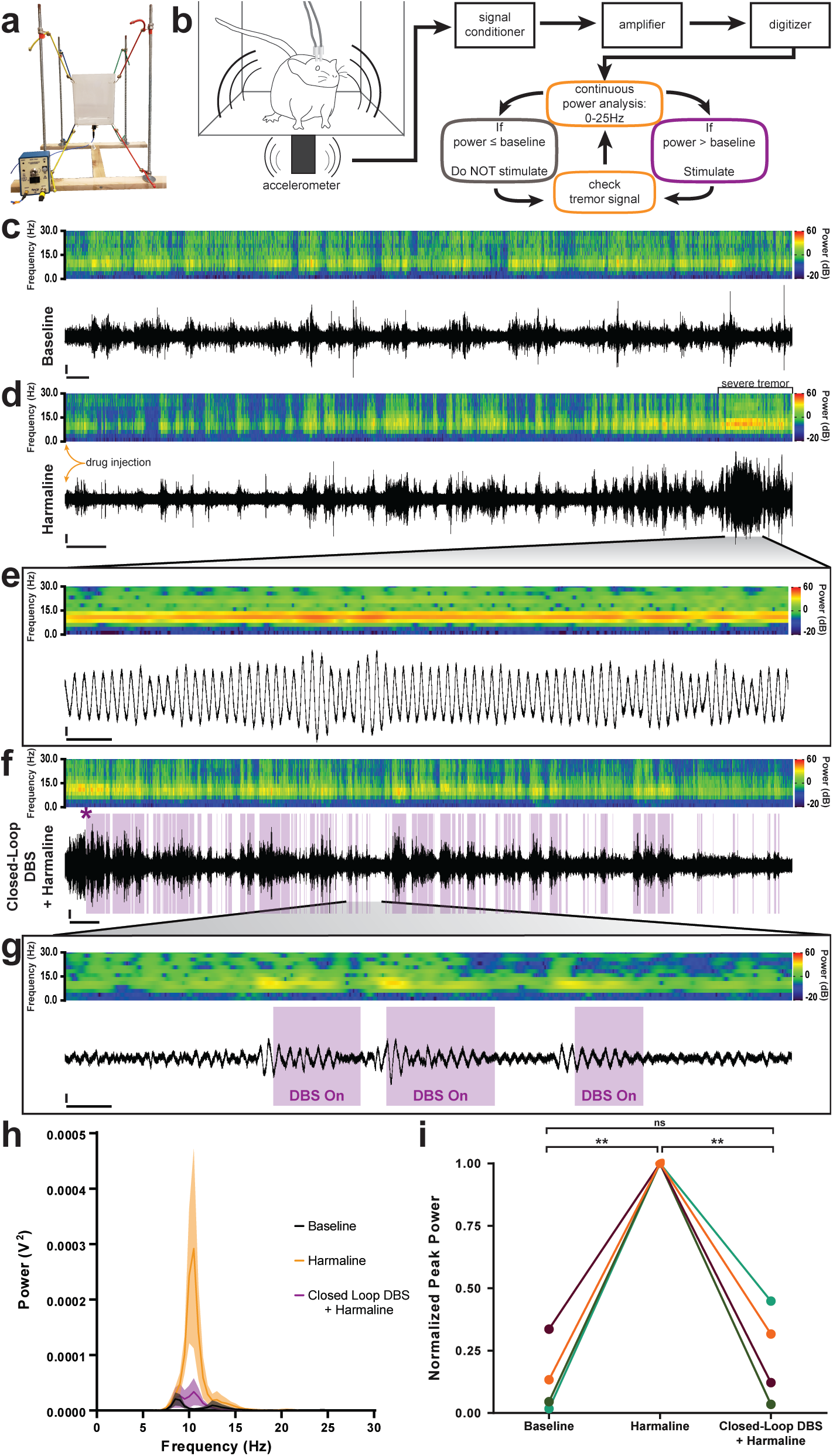
Closed-loop DBS targeted to the interposed nucleus reduces tremor severity. (**a**) Picture of the custom built tremor monitor. (**b**) Representation of the closed-loop design. Left: a mouse exhibiting tremor behavior shakes the tremor monitor chamber with an attached accelerometer (black rectangle). Accelerometer signals are passed through a signal conditioner, amplifier, and digitizer before a continuous power spectrum analysis is applied for frequencies from 0 Hz to 25 Hz. If tremor power exceeds a set threshold based on the individual’s baseline recording, stimulation is initiated for 36ms before re-evaluation of tremor power. No stimulation is generated if tremor power is below threshold level. (**c-g**) Example tremor traces from a single animal. Top: sonogram view (continuous power spectrum) of raw tremor trace. Heat scale = −20dB to 60dB. Bottom: raw tremor traces. Vertical scale = 25mV. (**c**) Baseline physiological tremor recording. Horizontal scale = 10s. (**d**) Tremor recording beginning directly after harmaline injection (yellow arrows). Severe tremor develops as shown at the right of the recording (black bracket). Horizontal scale = 50s. (**e**) Inset region from **d** (black lines) highlighting continuous, severe tremor. Horizontal scale = 0.5s. (**f**) Tremor recording beginning directly after **d** and just before closed-loop DBS protocol is initiated. Initiation time indicated by purple asterisk. Stimulation periods indicated by overlaid purple bars. Horizontal scale = 50s. (**g**) Inset region from **f** (black lines) highlighting bouts of tremor sufficient to cross threshold for stimulation. Stimulation periods indicated by overlaid purple bars. Horizontal scale = 0.5s. (**h**) Population average tremor traces for baseline, harmaline tremor, and closed-loop DBS during harmaline tremor periods. Solid line = average. Shaded region = SEM. N = 4. (**i**) Peak tremor power for each analyzed time period normalized to overall peak for all individuals. Power of tremor during harmaline period without closed-loop DBS stimulation was significantly greater than both baseline and the closed-loop DBS periods. There was no significant difference between baseline and the closed-loop DBS period. N = 4. Harmaline alone vs baseline: P = 0.0026. Harmaline alone vs closed-loop DBS + harmaline: P = 0.0077. Baseline vs closed-loop DBS + harmaline: P = 0.7772.

## Discussion

The participation of cerebellar dysfunction in a wide range of tremor disorders is universally anticipated (*10*). Here, we show that cerebellar neurons can produce the neural signals and behavioral states that are indicative of tremor. We demonstrate that it is not a lack of Purkinje cell activity, but instead an abnormal pattern of cerebellar output firing that causes tremor. Further, closed-loop DBS targeted to the cerebellar nuclei is sufficient to reduce pathological tremor.

While a loss of Purkinje cells and their ability to communicate with their downstream partners has been found in tremor disorders (*17*), we suggest here that pathological Purkinje cell activity – whether before, in the context of, or without cell death – is a strong impetus of tremor. We first showed evidence for this using *L7^Cre^;Vgat^flox/flox^* animals to demonstrate that Purkinje cell neurotransmission contributes to baseline physiological tremor. We further demonstrated that the lack of Purkinje cell neurotransmission abolishes harmaline tremor. The *L7^Cre^;Vgat^flox/flox^* approach used here is a constitutive genetic silencing model with high efficiency that causes no known gross morphological defects in any cerebellar cell type or degeneration (*27*). However, it is possible that eliminating Purkinje cell activity could result in subtle compensatory changes that suppress tremor (*27*). We note that some studies of human cases of tremor have found Purkinje cell loss as well as Purkinje cell degeneration and abnormal morphology (*17*). As our model is constitutive and *Vgat* is removed from Purkinje cells throughout the cerebellum, it does not mimic the progressive loss of Purkinje cells or a localized insult to cerebellar circuitry that has been associated with some manifestations of tremor in humans (*36, 37*). However, Purkinje cell loss is not always necessary for tremor to occur (*11, 23*). Therefore, because there are many forms of the disease, there could be many different potential mechanisms of tremor induction that do not involve Purkinje cell loss or degeneration (*11*). In cases of tremor where Purkinje cell loss is reported, the loss is incomplete and leaves to question what signals the remaining Purkinje cells send and how their downstream partners in the cerebellar nuclei respond (*24*). Indeed, cases of cerebellar stroke have been found to reduce tremor in humans (*38*). Thus, rather than questioning the anatomic or genetic changes associated with specific forms of tremor, our experiments argue that there may be a commonality of abnormal electrical signaling patterns that result in tremor in general. In this study, we have shown that a single cell type, the Purkinje cells, have the ability to serve as a gatekeeper of tremor.

We have found that if cerebellar nuclei cells are induced into bursting patterns of activity, a tremor will result. We show this pattern of activity of the cerebellar nuclei requires intact Purkinje cell output in the case of harmaline tremor, which has been used to model tremor behavioral phenotypes for the goal of developing therapeutics (*28*). Abnormal activity in the Purkinje cell to cerebellar nuclei connection is also implicated in ataxia (*39, 40*) and dystonia (*41–44*). This is intriguing because it raises the hypothesis that different modes of abnormal activity could contribute to ataxia, dystonia, and tremor. For example, in dystonia the cerebellar nuclei cells are induced into an irregular firing pattern with both elevated CV and CV2 (*44*) while in the electrophysiological recordings during tremor that we describe here show only an elevated CV. Instead of the erratic pattern of activity seen in dystonia, we observe a regular bursting pattern of activity in tremor. It is tempting to speculate that the spike firing plus population features such as synchrony (*45*) may distinguish these and other disease states that arise from or involve cerebellar circuitry. This idea is further supported by our data which demonstrates that multiple frequencies of cerebellar optogenetic stimulation can cause a wide range of tremor phenotypes, which in humans would equate to very different disease conditions. Our finding of optogenetically-induced tremor severity peaking at around 10 Hz matches the frequency of tremor commonly noted in humans with essential tremor (*3*) while the 3 Hz peak matches that frequently observed in Holmes’ tremor, which has suspected cerebellar involvement (*46*). The peak at 20 Hz is intriguing as it is both a harmonic of 10 Hz and similar frequencies have been described in genetic mouse models of tremor (*47*). We therefore postulate that, whether directly driven by abnormal Purkinje cell activity or other genetic, pharmacologic, and pathologic factors that influence cerebellar circuitry, any instigator of a synchronous, regular bursting pattern of cerebellar nuclei activity could result in tremor. Together, our data provide experimental support that selective alterations in cerebellar function are capable of producing a tremor phenotype across a range of disease-relevant frequencies. The data also underscore the capacity of functional heterogeneity, notably the heterogeneity in the defects arising from a common Purkinje cell circuit, that may promote the cerebellum’s involvement in multiple diseases. It is this inherent circuit flexibility that may also equip the cerebellum to contribute to a vast number of motor as well as non-motor behaviors.

Finally, our ability to acutely disrupt ongoing tremor behavior using closed-loop DBS suggests that the cerebellum itself may be an ideal target for the treatment of intractable tremor. This is consistent with the current practice of targeting DBS electrodes to regions of the thalamus that receive cerebellar input (*48*). Our previous work showed that DBS directed to the cerebellar nuclei ameliorates dystonia in mice (*44*). The addition of tremor as another hyperkinetic movement disorder that can be treated with cerebellar DBS provides hope that other cerebellum-associated movement disorders may be addressable with cerebellar DBS. Indeed, cerebellar DBS may be especially useful with patients who present with multiple cerebellum-related features, such as tremor with ataxia or tremor with dystonia (*49*). Moreover, the need for alternative targets has become apparent as some patients develop tolerance to thalamic DBS (*50*) or exhibit cognitive and motor decline as a result of current stimulation practices (*51, 52*). Importantly, in humans, targeting the cerebellar nuclei with neuromodulation has shown great promise after cerebellar stroke (*53*) and specifically for DBS, functional improvements were reported after dentate-directed DBS in rat models of cortical stroke (*54, 55*). While our understanding of the therapeutic mechanisms of DBS remains incomplete, both the data we present in this study and our previous work suggest that the disruption of abnormal patterns of Purkinje cell neurotransmission to the cerebellar nuclei and/or correction of abnormal cerebellar nuclei activity is instrumental to the success of cerebellar DBS. We show here that an abnormal pattern of activity that is transmitted from the Purkinje cells to the nuclei cells can be interrupted for recovery from tremor. In all, our data implicate a cerebellar circuit mechanism of tremor that may operate across tremor disorders and highlight the cerebellum as a potential target for tremor therapy.

## Acknowledgments

### Funding

This work was supported by Baylor College of Medicine, Texas Children’s Hospital, the National Institute of Neurological Disorders and Stroke (AMB: F31NS101891, JJW: F31NS092264, RVS: R01NS089664 and R01NS100874), the Eunice Kennedy Shriver National Institute of Child Health and Human Development (U54HD083092), a BCM IDDRC Project Development Award, the Hamill Foundation, the Mrs. Clifford Elder White Graham Endowed Research Fund, and the Bachmann-Strauss Dystonia and Parkinson Foundation, Inc. The content is solely the responsibility of the authors and does not necessarily represent the official views of the National Center for Research Resources or the National Institutes of Health.

## Materials and Methods

### Mouse lines

All experiments were performed according to a protocol approved by the Institutional Animal Care Use Committee (IACUC) of Baylor College of Medicine. Both male and female adult mice, at least two months of age, were studied. All mice were kept on a 14h/10h light/dark cycle. Purkinje cell specificity was achieved using a *L7^Cre^* transgenic mouse line (*25*). Genetic removal of Purkinje cell GABA neurotransmission was achieved by crossing this line to one that expresses a knock-in floxed *Vgat* allele (*26*). Optogenetic manipulation of Purkinje cells was achieved by crossing the *L7^Cre^* line to a reporter line that expresses channelrhodopsin (ChR2) fused to enhanced yellow fluorescent protein (EYFP) behind a *floxed-stop* cassette that was targeted to the *Rosa26* locus (*56*). During breeding, we considered the day a vaginal plug was visible as E0.5 and the day of birth as P0. We used standard genotyping protocols and primers for *Cre* and *Gfp* (to detect *Eyfp*) as described previously (*27, 44*) and the *Vgat floxed* allele detected as originally published (*26*).

### Tremor recording and analysis

Tremor was detected using at least one of three methods, including two tremor monitors and EMG. Tremor monitors used included a commercial model (San Diego Instruments, San Diego, CA, USA) as well as a custom built model that was inspired by a previously used design (*30*). In the commercial model setup, mice are placed inside a clear plastic tube that is fused to a small platform with rounded legs. An accelerometer is mounted to the bottom of the platform beneath the mouse and detects the shaking of the platform caused by the mouse’s tremor. The entire setup is placed inside of a sound-reducing opaque box. In the custom-built model, mice are placed into a translucent plastic box with an open top. The box is held steady in air by eight elastic cords, one end of each cord is connected to a separate corner of the box while the opposite ends are connected to vertical metal rods that form a perimeter around the box. The elastic cords that are connected to the top corners of the box are attached to the top of the nearest metal rod while the elastic cords that are connected to the bottom corners of the box are attached to the bottom of the nearest metal rod. The top cords provide upwards tension while the bottom cords provide downwards tension.

Mice were allowed to acclimate to the tremor monitor for at least 128 seconds before recordings of tremor were made. For both EMG and tremor monitor recordings, power spectrums of tremor were made using a fast Fourier transform (FFT) with a Hanning window. An offset was applied if the tremor waveform was not centered on 0 and the recordings were downsampled using the Spike2 software “interpolate” channel process in order to produce frequency bins aligned to whole numbers. FFT frequency resolution was targeted to either ∼0.25 Hz or ∼0.5 Hz per bin. Sonogram plots of tremor were made using the Spike2 software “sonogram” channel draw mode with a Hanning window. Power of tremor within a band was calculated by summing the power of each frequency within the band. Alpha + beta band was considered to be 8 Hz to 19.5 Hz. Gamma band was considered to be 20 Hz to 30 Hz. Brown-Forsythe and Welch ANOVA tests with Dunnett’s T3 multiple comparisons test were performed to determine whether band power was significantly different between conditions. For normalized peak comparison, an RM one-way ANOVA with Geisser-Greenhouse correction and Tukey’s multiple comparison’s test were used. A minimum of 25 seconds was analyzed for each animal in each time period.

Number of animals tested is represented by “N”. “Control” refers to *Vgat^flox/flox^* animals while “mutant” refers to *L7^Cre^;Vgat^flox/flox^* animals. P value > 0.05 = ns, ≤ 0.05 = *, ≤ 0.01 = **, ≤ 0.001 = ***, < 0.0001 = ****.

### Administration of harmaline

Adult mice were administered 30mg/kg harmaline (Sigma-Aldrich, St. Louis, MO, USA; #H1392) via intraperitoneal injection (IP). Harmaline tremor consistently developed between 5 to 15 minutes after injection. Mice were administered only one dose of harmaline and were sacrificed within 4 hours of the injection.

### Immunohistochemistry

Perfusion and tissue fixation were performed as previously described (*57*). In short, mice were anesthetized with Avertin (2, 2, 2-Tribromoethanol, Sigma-Aldrich, St. Louis, MO, USA; #T48402) via intraperitoneal injection. Once mice were deeply anesthetized, a whole-body perfusion was performed first with 0.1M phosphate-buffered saline (PBS; pH 7.4), then with 4% paraformaldehyde (4% PFA) diluted in PBS. The brain was then dissected out and placed in 4% PFA for 24 to 48 hours for post-fixation. Cryoprotection was then performed by placing the tissue in stepwise sucrose dilutions, first in 15% sucrose in PBS followed by 30% sucrose in PBS. After cryoprotection, the tissue was embedded in Tissue-Tek^®^ O.C.T. Compound (Sakura, Torrance, CA, USA) and frozen at −80°C. Tissue sections were then cut on a cryostat at 40µm thickness and stored free-floating in PBS at 4°C. Immunohistochemistry procedures have been described previously (*27, 58–60*). C-Fos staining was performed using rabbit polyclonal anti-c-Fos (Santa Cruz Biotechnology, Dallas, TX, USA; #sc-52). Signal was amplified using a Vectastain ABC kit (Vector Laboratories, Burlingame, CA, USA; #PK-6100) and followed with biotinylated goat anti-rabbit antibodies (Vector Laboratories, Burlingame, CA, USA; #BA-1000). Finally a 3, 3′-diaminobenzidine (DAB; Sigma-Aldrich, St Louis, MO, USA; #D5905-50TAB) reaction was used to reveal the antibody binding. After staining, sections were mounted on electrostatically coated slides with mounting medium (Vector Laboratories, Burlingame, CA, USA; #H-1200, Electron Microscopy Sciences, Hatfield, PA, USA; #17985-11 or Thermo Fisher Scientific, Waltham, MA, USA; #8312-4) and imaged. Triple fluorescent staining was completed using rabbit polyclonal anti-carbonic anhydrase VIII (Car8) to visualize Purkinje cells (CAVIII, Santa Cruz Biotechnology, #sc-67330), chicken anti-GFP to visualize ChR2 (Abcam, Cambridge, UK, #AB13970), and NeuroTrace fluorescent Nissl 530/615 to visualize neurons (Molecular Probes Inc., Eugene, OR, USA, #N21482). Secondary antibodies included Alexa-488 and -647 secondary goat anti-mouse and anti-rabbit antibodies (Molecular Probes Inc., Eugene, OR, USA).

### Imaging of immunostained tissue sections

Photomicrographs of stained tissue sections were captured using either a Zeiss Axio Imager.M2 microscope equipped with Zeiss AxioCam MRm and MRc5 cameras (Zeiss, Oberkochen, Germany) or a Leica DM4000 B LED microscope equipped with Leica DFC365 FX and Leica DMC 2900 cameras (Leica Microsystems Inc., Wetzlar, Germany). Zeiss Zen software was used for image acquisition from the Zeiss microscope. Leica Application Suite X (LAS X) software was used for image acquisition from the Leica microscope. Images were corrected for brightness and contrast using Adobe Photoshop CS5 (Adobe Systems, San Jose, CA, USA) for figure preparation. Schematics were made in Adobe Illustrator CC.

### Surgery

We have previously described our general surgical techniques in detail (*61*). In brief, for all surgical techniques used in these studies, mice were given preemptive analgesics (buprenorphine, 0.6mg/kg subcutaneous (SC), and meloxicam, 4mg/kg SC) with continued applications provided as part of the post-operative procedures. Anesthesia was induced with 3% isoflurane gas and maintained during surgery at 2% isoflurane gas. All surgeries were performed on a stereotaxic platform (David Kopf Instruments, Tujenga, CA, USA) with sterile surgery techniques. The following surgical techniques were either employed as individual experiments or combined depending on the requirements of the experiment:

#### Awake head-fixed neural recordings

During surgeries for awake neural recording experiments, the dorsal aspect of the skull was exposed and a circular craniotomy of about 2mm in diameter was performed dorsal to the interposed nucleus (6.4mm posterior and ±1.3mm lateral to Bregma.) A custom-built 3D-printed chamber was placed around the craniotomy and filled with antibiotic ointment. A custom headplate used to stabilize the mouse’s head during recordings was affixed over Bregma, and a skull screw was secured into an unused region of skull. All implanted items were secured using C&B Metabond Adhesive Luting Cement (Parkell, Edgewood, NY, USA) followed by a layer of dental cement (A-M Systems, Sequim, WA, USA; dental cement powder #525000 and solvent #526000) to completely enclose the area.

#### Optical fiber implantation

Optical fiber implant surgeries began as described above, however instead of performing a single large craniotomy, two small craniotomies were performed bilaterally and dorsal to the interposed nucleus through which two optical fibers (Thorlabs, Newton, NJ, USA; #FT200UMT) were lowered just into the region of the interposed nucleus. Optical fibers had been previously glued into ceramic ferrules (Thorlabs, Newton, NJ, USA; #CFLC230-10), polished (Thorlabs, Newton, NJ, USA; #LF5P, #LF3P, #LF1P, #LF03P, #LFCF), epoxied to each other at a set distance, and placed inside ceramic mating sleeves (Thorlabs, Newton, NJ, USA; #ADAL 1-5) prior to implantation. The fibers, a skull screw, and 1 or 2 metal rods used for holding the mouse’s head stable while connecting the optical fiber patch cables were affixed to the skull using C&B Metabond Adhesive Luting Cement followed by dental cement.

#### DBS electrode implantation

DBS electrode implant surgeries were performed exactly as the optical fiber implantation surgeries. Except, instead of optical fibers, custom 50 mm twisted bipolar Tungsten DBS electrodes were used (PlasticsOne, Roanoke, VA, USA; #8IMS303T3B01).

#### EMG implantation

Surgeries for EMG electrodes required only exposing the skull before an incision was made into the left hind limb. Handmade silver wire electrodes (A-M Systems, Sequim, WA, USA; #785500) were then implanted into the gastrocnemius of the left hind limb. The electrode wire was fed under the skin from the hind limb to the skull. An additional ground electrode was implanted under the skin of the neck. These wires were soldered to a connector for a detachable preamplifier (Pinnacle Technology, Inc., Lawrence, KS, USA; #8406). The connector, a skull screw, and 1 or 2 metal rods used for holding the mouse’s head stable while connecting the preamplifier were affixed to the skull using C&B Metabond Adhesive Luting Cement followed by dental cement.

### *In vivo* electrophysiology

Single-unit extracellular recordings were performed as previously described (*23, 44, 62*). Mice were awake and head-fixed to a frame while standing on a foam wheel which reduced the force they were able to apply to the headplate. Before recordings, mice were trained to become accustomed to being in the recording setup and head-fixed. At the time of recordings, the chamber around the craniotomy was emptied of antibiotic ointment and refilled with 0.9% w/v NaCl solution. Electrodes with 4-13MΩ impedance made of either tungsten (Thomas Recording, Giessen, Germany) or glass (Harvard Apparatus, Cambridge, MA, USA; #30-0057) which had been pulled at the time of recording (Sutter Instrument, Novato, CA, USA; #P-1000) were connected to a preamplifier headstage (NPI Electronic Instruments, Tamm, Germany). The headstage was attached to a motorized micromanipulator (MP-225; Sutter Instrument Co., Novato, CA, USA). Headstage output was amplified and bandpass filtered at 0.3-13 kHz (ELC-03XS amplifier, NPI Electronic Instruments, Tamm, Germany) before being digitized (CED Power 1401, CED, Cambridge, UK), recorded, and analyzed using Spike2 software (CED, Cambridge, UK).

Purkinje cells were identified by their location, firing rate, and presence of both complex and simple spike activity. Cerebellar nuclei cells were identified by their location and firing rate. Stable, clear, and continuous single-unit recordings of at least 30 seconds duration were included in the analysis. Firing properties were analyzed using Spike2 (CED, Cambridge, UK), Microsoft Excel (Microsoft, Redmond, WA, USA), custom MATLAB code (MathWorks, Natick, MA, USA), and GraphPad Prism (GraphPad Software, La Jolla, CA, USA) software. The number of animals tested is represented by “N” while the number of cells recorded is represented by “n”. “Control” refers to *Vgat^flox/flox^* animals while “mutant” refers to *L7^Cre^;Vgat^flox/flox^* animals. P value > 0.05 = ns, ≤ 0.05 = *, ≤ 0.01 = **, ≤ 0.001 = ***, < 0.0001 = ****.

### Optogenetic Stimulation

Optical fibers were either implanted into the region of the interposed nucleus (see above) or held close to the tip of a recording electrode via an optopatcher (ALA Scientific Instruments Inc., Farmingdale, NY, USA) or a custom-built electrode setup. Stimulation patterns were programmed and recorded using Spike2 software and delivered using a CED Power1401 data acquisition interface (CED, Cambridge, UK) to control a 465nm LED (ALA Scientific Instruments Inc., Farmingdale, NY, USA). Maximum LED power at the end of the fiber patch cable was measured to be 3.6mW and stimulation consisted of sinusoidal pulses of light, from light off to maximum brightness to light off.

### Closed-Loop DBS

Closed-loop DBS programs were written as custom Spike2 scripts and configuration files (CED, Cambridge, UK). To stimulate in the condition of tremor, we wrote scripts that ran in the background of our recordings and functioned in concert with the configuration paradigm that acted as the stimulation generator. The scripts were designed to detect and trigger DBS based on the presence of an enhanced tremor phenotype.

To detect tremor behavior, tremor monitor activity between 0 and 25 Hz was used to prompt either the start or stop DBS stimulation depending on whether this abnormal activity was present. First our script created an analysis channel in our recordings that continuously performed a power analysis of the 0 to 25 Hz frequency range of our raw tremor monitor signal using a Fast Fourier Transform (FFT) with a resolution of 0.6104 Hz, size of 8192 points, and sampled over a period of 1.6384 seconds. The script sampled the output of this analysis as close to real time as possible to determine whether the power of the band had surpassed a threshold that had been set to the individual mouse’s peak tremor power from the baseline recording. If the threshold had been surpassed, a start signal was generated.

The start signal prompted a simultaneously-running Spike2 configuration paradigm to initiate DBS. The program initiated a looping pulse train of 5V pulses of 20μs duration to be output from a Power1401 (CED, Cambridge, UK). The pulses occurred with an interval of 8ms and began 1ms after the start signal was received. This produced stimulation at 125 Hz. The pulses were sent to a Master-8 stimulator (A.M.P.I., Jerusalem, Israel) that allowed precision timing in the production of 60μs square biphasic pulses. These pulses were then output to external stimulus isolators (ISO-Flex, A.M.P.I., Jerusalem, Israel) that set the DBS current to 30µA. The pulse train lasted at minimum 36ms. Within the final 8ms gap between pulses of the loop, the program would check for the start signal. If the start signal was still present (i.e. tremor was ongoing), the pulse train would continue for another 36ms. If the start signal was no longer present (i.e. tremor had returned to below threshold levels), DBS stimulation would halt until the start signal was generated again. Post hoc analysis of tremor was performed using Spike2 software and custom Matlab scripts (MathWorks, Natick, MA, USA).

## Movies

**Movie S1:** *L7^Cre^;Vgat^flox/flox^* mice exhibit ataxia and disequilibrium, but not pathological tremor.

**Movie S2:** Harmaline induces a severe tremor in *Vgat^flox/flox^* control mice but not in *L7^Cre^;Vgat^flox/flox^* mutant mice.

**Movie S3:** Optogenetic stimulation of Purkinje cell terminals in the interposed nuclei of *L7^Cre^;ROSA26^flox-stop-ChR2-EYFP^* mice using a 10 Hz sinusoidal stimulus induces a robust tremor.

