## Supplementary material for "Cerebellar Purkinje cell microcircuits are essential for tremor": Table 1

| Figure | Comparator 1 | Comparator 2 | Mean 1 | Mean 2 | n 1 | n 2 | Summary | Adjusted P Value |
|---|---|---|---|---|---|---|---|---|
| Fig. 1h | control baseline | control + harmaline | 0.002181 $V^2$ | 0.02005 $V^2$ | 16 | 16 | * | 0.0190 |
| | control baseline | mutant baseline | 0.002181 $V^2$ | 0.001438 $V^2$ | 16 | 12 | ns | 0.2982 |
| | control baseline | mutant + harmaline | 0.002181 $V^2$ | 0.002787 $V^2$ | 16 | 13 | ns | 0.9420 |
| | control + harmaline | mutant baseline | 0.02005 $V^2$ | 0.001438 $V^2$ | 16 | 12 | * | 0.0142 |
| | control + harmaline | mutant + harmaline | 0.02005 $V^2$ | 0.002787 $V^2$ | 16 | 13 | * | 0.0252 |
| | mutant baseline | mutant + harmaline | 0.001438 $V^2$ | 0.002787 $V^2$ | 12 | 13 | ns | 0.3550 |
| Fig. 1i | control baseline | control + harmaline | 0.001032 $V^2$ | 0.001767 $V^2$ | 16 | 16 | ns | 0.2224 |
| | control baseline | mutant baseline | 0.001032 $V^2$ | 0.0005844 $V^2$ | 16 | 12 | * | 0.0454 |
| | control baseline | mutant + harmaline | 0.001032 $V^2$ | 0.0008114 $V^2$ | 16 | 13 | ns | 0.8693 |
| | control + harmaline | mutant baseline | 0.001767 $V^2$ | 0.0005844 $V^2$ | 16 | 12 | * | 0.0155 |
| | control + harmaline | mutant + harmaline | 0.001767 $V^2$ | 0.0008114 $V^2$ | 16 | 13 | ns | 0.0897 |
| | mutant baseline | mutant + harmaline | 0.0005844 $V^2$ | 0.0008114 $V^2$ | 12 | 13 | ns | 0.8543 |
