## Supplementary material for "Cerebellar Purkinje cell microcircuits are essential for tremor": Table 2

| Figure | Comparator 1 | Comparator 2 | Mean 1 | Mean 2 | n 1 | n 2 | Summary | Adjusted P Value |
|---|---|---|---|---|---|---|---|---|
| Fig. 2h | control baseline | control + harmaline | 77.63 Hz | 49.18 Hz | 18 | 14 | ** | 0.0020 |
| | control baseline | mutant baseline | 77.63 Hz | 55.97 Hz | 18 | 15 | * | 0.0238 |
| | control baseline | mutant + harmaline | 77.63 Hz | 26.65 Hz | 18 | 12 | **** | <0.0001 |
| | control + harmaline | mutant baseline | 49.18 Hz | 55.97 Hz | 14 | 15 | ns | 0.8208 |
| | control + harmaline | mutant + harmaline | 49.18 Hz | 26.65 Hz | 14 | 12 | * | 0.0419 |
| | mutant baseline | mutant + harmaline | 55.97 Hz | 26.65 Hz | 15 | 12 | ** | 0.0037 |
| Fig. 2i | control baseline | control + harmaline | 0.4902 | 1.372 | 18 | 14 | *** | 0.0003 |
| | control baseline | mutant baseline | 0.4902 | 1.152 | 18 | 15 | ** | 0.0079 |
| | control baseline | mutant + harmaline | 0.4902 | 1.953 | 18 | 12 | **** | <0.0001 |
| | control + harmaline | mutant baseline | 1.372 | 1.152 | 14 | 15 | ns | 0.7229 |
| | control + harmaline | mutant + harmaline | 1.372 | 1.953 | 14 | 12 | ns | 0.0552 |
| | mutant baseline | mutant + harmaline | 1.152 | 1.953 | 15 | 12 | ** | 0.0032 |
| Fig. 2j | control baseline | control + harmaline | 0.4443 | 0.6683 | 18 | 14 | *** | 0.0005 |
| | control baseline | mutant baseline | 0.4443 | 0.5355 | 18 | 15 | ns | 0.3088 |
| | control baseline | mutant + harmaline | 0.4443 | 0.8396 | 18 | 12 | **** | <0.0001 |
| | control + harmaline | mutant baseline | 0.6683 | 0.5355 | 14 | 15 | ns | 0.0897 |
| | control + harmaline | mutant + harmaline | 0.6683 | 0.8396 | 14 | 12 | * | 0.0253 |
| | mutant baseline | mutant + harmaline | 0.5355 | 0.8396 | 15 | 12 | **** | <0.0001 |
| Fig. 2k | control baseline | control + harmaline | 1.289 Hz | 2.876 Hz | 18 | 14 | *** | 0.0003 |
| | control baseline | mutant baseline | 1.289 Hz | 1.458 Hz | 18 | 15 | ns | 0.9649 |
| | control baseline | mutant + harmaline | 1.289 Hz | 5.338 Hz | 18 | 12 | **** | <0.0001 |
| | control + harmaline | mutant baseline | 2.876 Hz | 1.458 Hz | 14 | 15 | ** | 0.0025 |
| | control + harmaline | mutant + harmaline | 2.876 Hz | 5.338 Hz | 14 | 12 | **** | <0.0001 |
| | mutant baseline | mutant + harmaline | 1.458 Hz | 5.338 Hz | 15 | 12 | **** | <0.0001 |
| Fig. 2l | control baseline | control + harmaline | 0.7396 | 0.5753 | 18 | 14 | ns | 0.1609 |
| | control baseline | mutant baseline | 0.7396 | 0.8631 | 18 | 15 | ns | 0.3751 |

| | | | | | | | | |
|---|---|---|---|---|---|---|---|---|
| | control<br>baseline | mutant<br>+ harmaline | 0.7396 | 0.4335 | 18 | 12 | ** | 0.0022 |
| | control<br>+ harmaline | mutant<br>baseline | 0.5753 | 0.8631 | 14 | 15 | ** | 0.0043 |
| | control<br>+ harmaline | mutant<br>+ harmaline | 0.5753 | 0.4335 | 14 | 12 | ns | 0.3572 |
| | mutant<br>baseline | mutant<br>+ harmaline | 0.8631 | 0.4335 | 15 | 12 | **** | <0.0001 |
| Fig. 2m | control<br>baseline | control<br>+ harmaline | 0.8876 | 0.5399 | 18 | 14 | **** | <0.0001 |
| | control<br>baseline | mutant<br>baseline | 0.8876 | 0.9082 | 18 | 15 | ns | 0.9848 |
| | control<br>baseline | mutant<br>+ harmaline | 0.8876 | 0.3224 | 18 | 12 | **** | <0.0001 |
| | control<br>+ harmaline | mutant<br>baseline | 0.5399 | 0.9082 | 14 | 15 | **** | <0.0001 |
| | control<br>+ harmaline | mutant<br>+ harmaline | 0.5399 | 0.3224 | 14 | 12 | ** | 0.0085 |
| | mutant<br>baseline | mutant<br>+ harmaline | 0.9082 | 0.3224 | 15 | 12 | **** | <0.0001 |
| Fig. 2n | control<br>baseline | control<br>+ harmaline | 0.008528 s | 0.04651 s | 18 | 14 | ** | 0.0015 |
| | control<br>baseline | mutant<br>baseline | 0.008528 s | 0.02349 s | 18 | 15 | ns | 0.4089 |
| | control<br>baseline | mutant<br>+ harmaline | 0.008528 s | 0.09622 s | 18 | 12 | **** | <0.0001 |
| | control<br>+ harmaline | mutant<br>baseline | 0.04651 s | 0.02349 s | 14 | 15 | ns | 0.1202 |
| | control<br>+ harmaline | mutant<br>+ harmaline | 0.04651 s | 0.09622 s | 14 | 12 | *** | 0.0001 |
| | mutant<br>baseline | mutant<br>+ harmaline | 0.02349 s | 0.09622 s | 15 | 12 | **** | <0.0001 |
| Fig. 2o | control<br>baseline | control<br>+ harmaline | 0.01806 s | 0.01806 s | 18 | 14 | ns | >0.9999 |
| | control<br>baseline | mutant<br>baseline | 0.01806 s | 0.03709 s | 18 | 15 | * | 0.0228 |
| | control<br>baseline | mutant<br>+ harmaline | 0.01806 s | 0.02643 s | 18 | 12 | ns | 0.6161 |
| | control<br>+ harmaline | mutant<br>baseline | 0.01806 s | 0.03709 s | 14 | 15 | * | 0.0358 |
| | control<br>+ harmaline | mutant<br>+ harmaline | 0.01806 s | 0.02643 s | 14 | 12 | ns | 0.6560 |
| | mutant<br>baseline | mutant<br>+ harmaline | 0.03709 s | 0.02643 s | 15 | 12 | ns | 0.4462 |
| Fig. 2p | control<br>baseline | control<br>+ harmaline | 59.78 | 17.71 | 18 | 17 | ** | 0.0035 |
| | control<br>baseline | mutant<br>baseline | 59.78 | 53.83 | 18 | 18 | ns | 0.9551 |
| | control<br>baseline | mutant<br>+ harmaline | 59.78 | 8.421 | 18 | 15 | *** | 0.0004 |
| | control<br>+ harmaline | mutant<br>baseline | 17.71 | 53.83 | 17 | 18 | * | 0.0155 |
| | control<br>+ harmaline | mutant<br>+ harmaline | 17.71 | 8.421 | 17 | 15 | ns | 0.8730 |

| | | | | | | | | |
|---|---|---|---|---|---|---|---|---|
| | mutant baseline | mutant + harmaline | 53.83 | 8.421 | 18 | 15 | ** | 0.0021 |
