## Supplementary material for "Cerebellar Purkinje cell microcircuits are essential for tremor": Table 3

| Figure | Comparator 1 | Comparator 2 | Mean 1 | Mean 2 | n 1 | n 2 | Summary | Adjusted P Value |
|---|---|---|---|---|---|---|---|---|
| Fig. 3f | control baseline | control + harmaline | 66.48 Hz | 74.56 Hz | 19 | 14 | ns | 0.8660 |
| | control baseline | mutant baseline | 66.48 Hz | 79.81 Hz | 19 | 18 | ns | 0.5246 |
| | control baseline | mutant + harmaline | 66.48 Hz | 82.22 Hz | 19 | 11 | ns | 0.5035 |
| | control + harmaline | mutant baseline | 74.56 Hz | 79.81 Hz | 14 | 18 | ns | 0.9593 |
| | control + harmaline | mutant + harmaline | 74.56 Hz | 82.22 Hz | 14 | 11 | ns | 0.9180 |
| | mutant baseline | mutant + harmaline | 79.81 Hz | 82.22 Hz | 18 | 11 | ns | 0.9966 |
| Fig. 3g | control baseline | control + harmaline | 0.4511 | 0.8654 | 19 | 14 | **** | <0.0001 |
| | control baseline | mutant baseline | 0.4511 | 0.3428 | 19 | 18 | ns | 0.5379 |
| | control baseline | mutant + harmaline | 0.4511 | 0.3695 | 19 | 11 | ns | 0.8150 |
| | control + harmaline | mutant baseline | 0.8654 | 0.3428 | 14 | 18 | **** | <0.0001 |
| | control + harmaline | mutant + harmaline | 0.8654 | 0.3695 | 14 | 11 | **** | <0.0001 |
| | mutant baseline | mutant + harmaline | 0.3428 | 0.3695 | 18 | 11 | ns | 0.9918 |
| Fig. 3h | control baseline | control + harmaline | 0.4141 | 0.5262 | 19 | 14 | ns | 0.0520 |
| | control baseline | mutant baseline | 0.4141 | 0.2310 | 19 | 18 | *** | 0.0001 |
| | control baseline | mutant + harmaline | 0.4141 | 0.2743 | 19 | 11 | * | 0.0179 |
| | control + harmaline | mutant baseline | 0.5262 | 0.2310 | 14 | 18 | **** | <0.0001 |
| | control + harmaline | mutant + harmaline | 0.5262 | 0.2743 | 14 | 11 | **** | <0.0001 |
| | mutant baseline | mutant + harmaline | 0.2310 | 0.2743 | 18 | 11 | ns | 0.7864 |
