## Supplementary material for "Cerebellar Purkinje cell microcircuits are essential for tremor": Table 4

| Figure | Comparator 1 | Comparator 2 | Mean 1 | Mean 2 | n 1 | n 2 | Summary | Adjusted P Value |
|---|---|---|---|---|---|---|---|---|
| Fig. 4n | pre | during | 0.1825 | 1.000 | 7 | 7 | **** | <0.0001 |
| | pre | post | 0.1825 | 0.1702 | 7 | 7 | ns | 0.9796 |
| | during | post | 1.000 | 0.1702 | 7 | 7 | *** | 0.0001 |
