## Supplementary material for "Cerebellar Purkinje cell microcircuits are essential for tremor": Table 5

| Figure | Comparator 1 | Comparator 2 | Mean 1 | Mean 2 | n 1 | n 2 | Summary | Adjusted P Value |
|---|---|---|---|---|---|---|---|---|
| Fig. 5i | baseline | harmaline | 0.1333 | 1.000 | 4 | 4 | ** | 0.0026 |
| | baseline | closed-loop DBS + harmaline | 0.1333 | 0.2305 | 4 | 4 | ns | 0.7772 |
| | harmaline | closed-loop DBS + harmaline | 1.000 | 0.2305 | 4 | 4 | ** | 0.0077 |
